# Infection with novel *Bacteroides phage BV01* alters host transcriptome and bile acid metabolism in a common human gut microbe

**DOI:** 10.1101/2020.04.06.028910

**Authors:** Danielle E. Campbell, Lindsey K. Ly, Jason M. Ridlon, Ansel Hsiao, Rachel J. Whitaker, Patrick H. Degnan

## Abstract

The bacterial genus *Bacteroides* is among the most abundant and common taxa in the human gut, yet little is known about the phages infecting the group. *Bacteroides phage BV01* (BV01) was identified as a prophage integrated on the chromosome of its host, *Bacteroides vulgatus* ATCC 8482. Phage BV01 is actively produced, and infects susceptible *B. vulgatus* hosts in the mouse gut. Infection with BV01 causes a generalized repression of the *B. vulgatus* transcriptome, downregulating 103 transcripts and upregulating only 12. Integration of BV01 disrupts the promoter sequence of a downstream gene encoding a putative tryptophan-rich sensory protein (*tspO*). Deletion of *tspO* and subsequent RNAseq analysis revealed that more than half of the differentially-regulated transcripts are shared with the BV01 lysogen, suggesting the transcriptomic response to BV01 is linked to *tspO*. Among these differentially-regulated transcripts are two encoding bile salt hydrolases. Bile acid deconjugation assays show that BV01 represses its host’s ability to hydrolyze bile acids in a *tspO*-dependent manner. Analysis of 256 published healthy human gut metagenomes suggests that phage integration adjacent to *B. vulgatus*-like *tspO* genes is rare within an individual, but common among humans. Finally, this work proposes a novel phage family that includes BV01, the *Salyersviridae*, whose host range spans the *Bacteroides* and is detectable in human-associated samples. Together, these findings highlight the importance of phage-host interactions to our understanding of how gut microbes sense and interact with their environment.

**IMPORTANCE:** The links between human disease and the gut microbiome are numerous. Most mechanisms by which most gut microbes and their activities change and impact human health remain elusive. Phages, viruses that infect bacteria, are hypothesized to play a central role in modulating both community dynamics and functional activities. Here we have characterized an active prophage, BV01, which infects a pervasive and abundant human gut-associated species. BV01 infection alters its host’s transcriptional profile including its metabolism of bile acids, molecules implicated in mediating health and disease states in the gut. This highlights that prophages and other components of the variable genome should not be overlooked in bacterial genomes because they may dramatically alter host phenotypes. Furthermore, BV01 represents a new family of phages infecting human gut symbionts, providing a foundation for future investigations of phage-host interactions in these clinically-relevant but underexplored hosts.

## INTRODUCTION

The human gut is colonized by a dense and diverse microbial community comprised of bacterial, archaeal, and fungal cells, as well as the viruses that infect them. This gut microbiome is vital to human health and development, and is linked to an increasingly long list of disease states. Recent work has specifically implicated the gut phageome in disease, including inflammatory bowel disease, malnutrition, AIDS, colorectal cancers, and hypertension (1–5). Broadly, gut phages act as important modulators of bacterial community structure (4, 6, 7) and metabolism (8). Despite their apparent importance, little is known about how most gut-associated phages interact with their bacterial hosts.

The *Bacteroides* is one of the most common and abundant bacterial genera in the distal human gut. The genus is known to degrade a diversity of complex carbohydrates (9, 10), and interact with host immune cells (11, 12). Within a single human host, many *Bacteroides* species and strains coexist, competing for nutrients under changing environmental conditions caused by host diet (13), host metabolites (14), host immune system activities (15, 16), and phage predation (17). Moreover, horizontal gene transfer plays an important role in shaping the evolution and function of *Bacteroides* genomes (18, 19). How the diversity of *Bacteroides* strains in the human gut persists over time in such a dense, dynamic, and competitive environment is likely multi-fold, and perhaps afforded by their highly plastic genomes.

Phage diversity and phage-host interactions within most commensal gut-associated bacteria, including *Bacteroides* species, is underexplored. Currently, the most abundant gut-associated phages are the crAssphages (20, 21), a group of related lytic phages that infect *Bacteroides intestinalis* and potentially other species. CrAssphages demonstrate how traditional phage techniques (*e.g.,* agar overlay plaque assays) are not reliable for *Bacteroides* hosts (22), likely due to heterogeneity in capsular polysaccharide composition within isogenic cultures (23, 24). In fact, deletion of all capsular polysaccharide synthesis loci allows for the isolation of many phages on the host *Bacteroides thetaiotaomicron* VPI-5482 (17). Most phages isolated against *Bacteroides* hosts thus far exhibit an obligately lytic lifestyle (17, 22, 25, 26), despite the potentially large role of lysogeny in phage-host interactions in the gut, where at least 17% of the gut phageome is predicted to be temperate (27, 28).

Prophage–host interactions have the potential to cause complex alterations to the host phenotype by virtue of the temperate lifestyle having two distinct phases: lysogeny and lysis. Unlike most strictly lytic phages, temperate phages may horizontally transfer beneficial genes between hosts, such as antibiotic resistance genes (29) and auxiliary metabolic genes (30, 31). Some phage regulatory machinery expressed from prophages can modulate the transcription of host genes, resulting in altered phenotypes (32, 33). Integration of prophages into the host genome may also disrupt or enhance the activity of surrounding chromosomal genes (34–36). While much is known about how these prophage–host interactions contribute to virulence in pathogens (37–41), exceedingly little is about how temperate phages modulate the activities of commensals.

Here we have identified an active prophage, *Bacteroides phage BV01,* in a genetically tractable host strain, *B. vulgatus* ATCC 8482, and characterized its effects on the host’s transcriptome and phenotype. Further we determine that BV01 represents a larger group of *Bacteroides-*associated prophages comprising the proposed phage family *Salyersviridae*, which are common members of the human gut phageome. This work provides the first insights to how the *Bacteroides* react to temperate phage infection, and establishes a model system for exploring complex phage–host interactions in an important human gut symbiont.

## RESULTS

### *Bacteroides phage BV01* is a prophage in *B. vulgatus* ATCC 8482

*Bacteroides phage BV01* was first partially predicted with ProPhinder and deposited in the ACLAME database (42, 43). Through comparative genomics and annotation of this region, we extended the predicted BV01 prophage to 58.9 Kb (NC_009614.1: 3,579,765..3,638,687), comprised of 72 predicted ORFs (Fig. 1), a prediction that agrees with an observation of prophage induction from *B. vulgatus* ATCC 8482 in a gnotobiotic mouse model (44). BV01 encodes genes suggesting a temperate lifestyle, with a putative CI-like repressor and a Cro-like anti-repressor, as well as a holing-lysin-spanin operon (Fig. 1).

**Figure 1.**
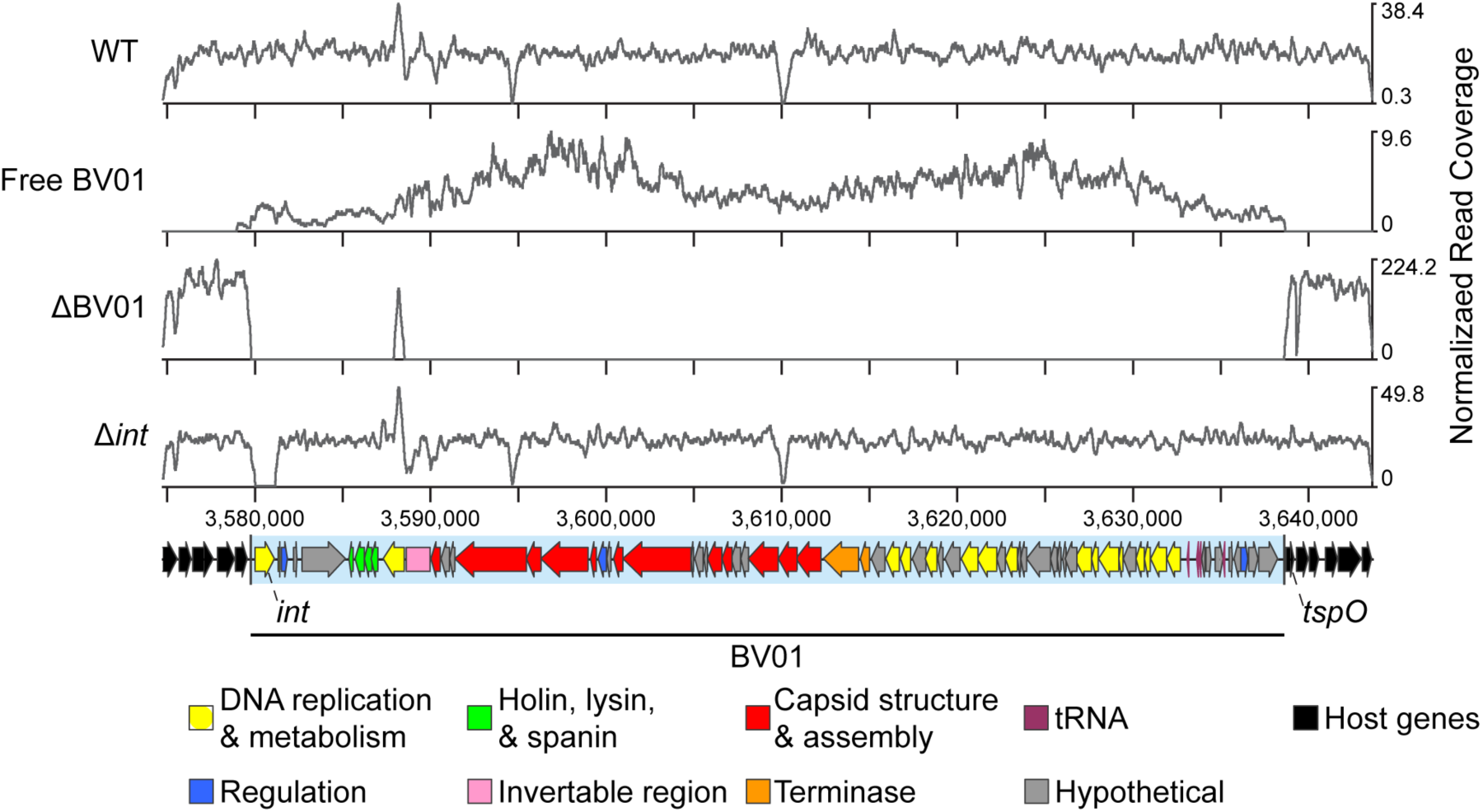
DNA sequencing reads mapped to the BV01 prophage region. Reads from shotgun sequencing of *B. vulgatus* genomic DNA (WT) or isolated phage DNA (Free BV01) were normalized to the total number of reads after trimming, and represented as a coverage curve. A cured lysogen (ΔBV01) and integrase deletion mutant (Δ*int*) of *B. vulgatus* were confirmed by shotgun sequencing of genomic DNA. The discrete coverage peak at position ∼3,588,000 nt from ΔBV01 is attributed to a homologous sequence elsewhere on the *B. vulgatus* chromosome. Putative functions of BV01 genes are indicated by the colors in the legend.

BV01 is detectable outside of host cells in the supernatants of *in vitro* cultures as a DNase-protected, dsDNA genome by both sequencing (Fig. 1) and PCR (Fig. 2A). An isogenic cured lysogen (ΔBV01) strain constructed by replacement with the corresponding chromosomal region of an uninfected *B. vulgatus* strain (*attB*) does not release free BV01 (Fig. 1, 2A). Assembly of sequencing reads from free BV01 phage DNA results in a circular contig that spans the phage attachment site (*attP*). The BV01 *attP* is identical to the left and right attachment sites (*attL* and *attR*), a pair of 25-bp direct repeat sequences (5’-GTCTAGTTTAGTTTTTGTGTTGTAA-3’), suggesting BV01 enters a circular intermediate before replication.

**Figure 2.**
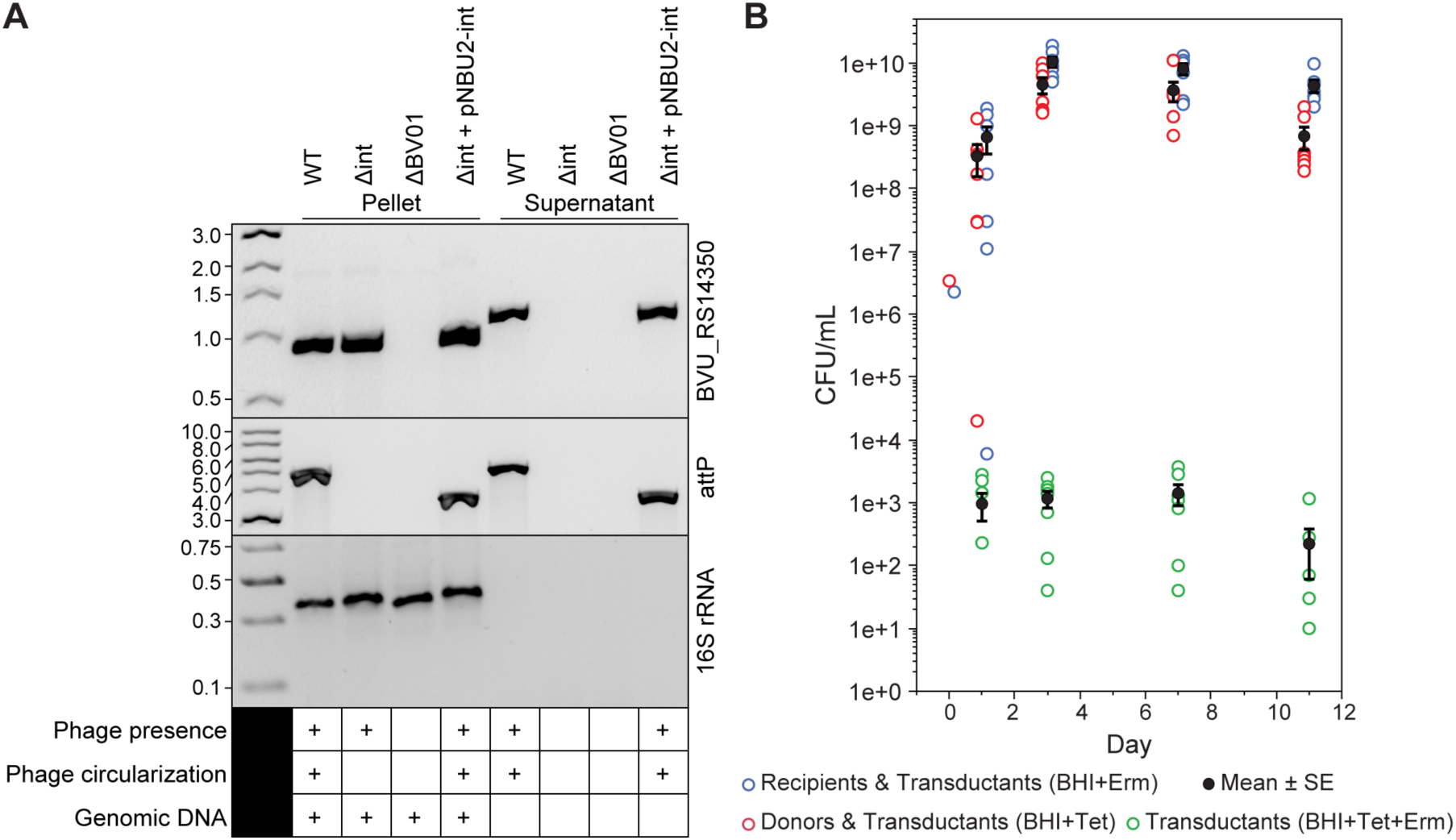
Prophage BV01 is an intact prophage. (A) Excision and circularization activities of the BV01 integrase are confirmed by PCR. The presence of phage DNA was detected by amplification of a phage marker gene (BVU_RS14350). Amplification across the phage attachment site (*attP*) indicates circularization of the BV01 genome; *attP* amplicons from the integrase complement strain (Δ*int* + pNBU2-*int*) are ∼1.2 Kb shorter than wild-type amplicons due to deletion of the integrase gene. Supernatant fractions were treated with DNase, eliminating all contaminating host genomic DNA, as demonstrated by the amplification of a host marker gene (16S rRNA). Note that despite apparent size shift of BVU_RS14350 amplicons from the pellets and supernatants, Sanger sequencing validated that the products are in fact identical. PCR amplicons were visualized by agarose gel electrophoresis alongside NEB 1 Kb DNA ladder (BVU_RS14350, *attP*) or GeneRuler Express DNA ladder (16S rRNA); ladder band sizes shown in Kb. (B) BV01 can transduce uninfected hosts in a gnotobiotic mouse. Germ-free mice (*n*=7) were gavaged with an equal mixture of a BV01-*tetQ* lysogen and an erythromycin-tagged cured lysogen (Day 0). Recipient, donor, and transductant cells were identified by plating on Brain Heart Infusion (BHI) media with antibiotic selection: erythromycin (Erm) or tetracycline (Tet).

To confirm that the release of DNase-protected BV01 genomes from host cells is a phage-encoded and directed process, we sought to identify the phage integrase. BV01 encodes three genes with integrase domains (PF00589); we hypothesized that the gene BVU_RS14130, adjacent to the phage attachment site, was the most likely candidate for catalyzing integration and excision of BV01. A BVU_RS14130 deletion mutant (Δ*int*) was constructed (Fig. 1) and its activity assayed by PCR of paired cell pellets and DNase-treated culture supernatants (Fig. 2A). Phage DNA was not detected in the supernatants of the Δ*int* strain. Furthermore, the Δ*int* mutant does not yield an amplicon for the circularization of the BV01. Expression of the *int* gene *in trans* from a pNBU2 plasmid complements both the circularization and release phenotypes (Fig. 2A). These results demonstrate that the integrase encoded by BVU_RS14130 is necessary for phage excision, circularization, and release from the host. Furthermore, they suggest that BV01 is an intact prophage capable of directing its own mobilization.

Despite numerous attempts, we have not identified *in vitro* inducing conditions for BV01. This has included treatments with UV light and sub-inhibitory concentrations of norfloxacin and mitomycin C (data not shown). BV01 supernatants never produced a plaque on any of 10 *B. vulgatus* isolates tested, the *B. vulgatus* cured lysogen, *B. thetaiotaomicron* VPI-5482, or *Bacteroides dorei* DSM 17855, on any of four media: TYG, BHI-HB, BHI-HM, and TYG_S_. To test for new lysogenic infections, BV01 was tagged with a copy of *tetQ*, conferring tetracycline resistance, using allelic exchange. Transduction of BV01 or BV01-*tetQ* via culture supernatants or in co-culture with erythromycin resistant hosts tagged with pNBU2-*bla*-*ermG*_b_ has never generated transductants. We conclude that BV01 is a latent prophage in culture, which is further supported by transcriptional data showing that BV01 exists in a largely repressed state (Fig. S1).

The only infectious conditions that have been identified for BV01 are in a gnotobiotic mouse model (Fig. 2B). Within a single day of gavage, BV01-*tetQ* transductants were identified on doubly-selective media from mouse pellets from 4 of the 7 mice. Over the course of the 11-day experiment transductants were eventually observed in all animals, in all cages. The average frequency of transduction ranged from 1.9 x 10^-6^ to 3.6 x 10^-9^ per animal. These results support the hypothesis that an unknown mammalian host factor is required for novel BV01 infection.

### Lysogeny with BV01 alters the host transcriptome

It was hypothesized that lysogeny with a prophage such as BV01 could alter the activities of the *B. vulgatus* host. RNA sequencing (RNAseq) of the *B. vulgatus* wild-type lysogen and the isogenic cured lysogeny was performed to identify transcripts differentially regulated in response to BV01 lysogeny (Fig. 3).

**Figure 3.**
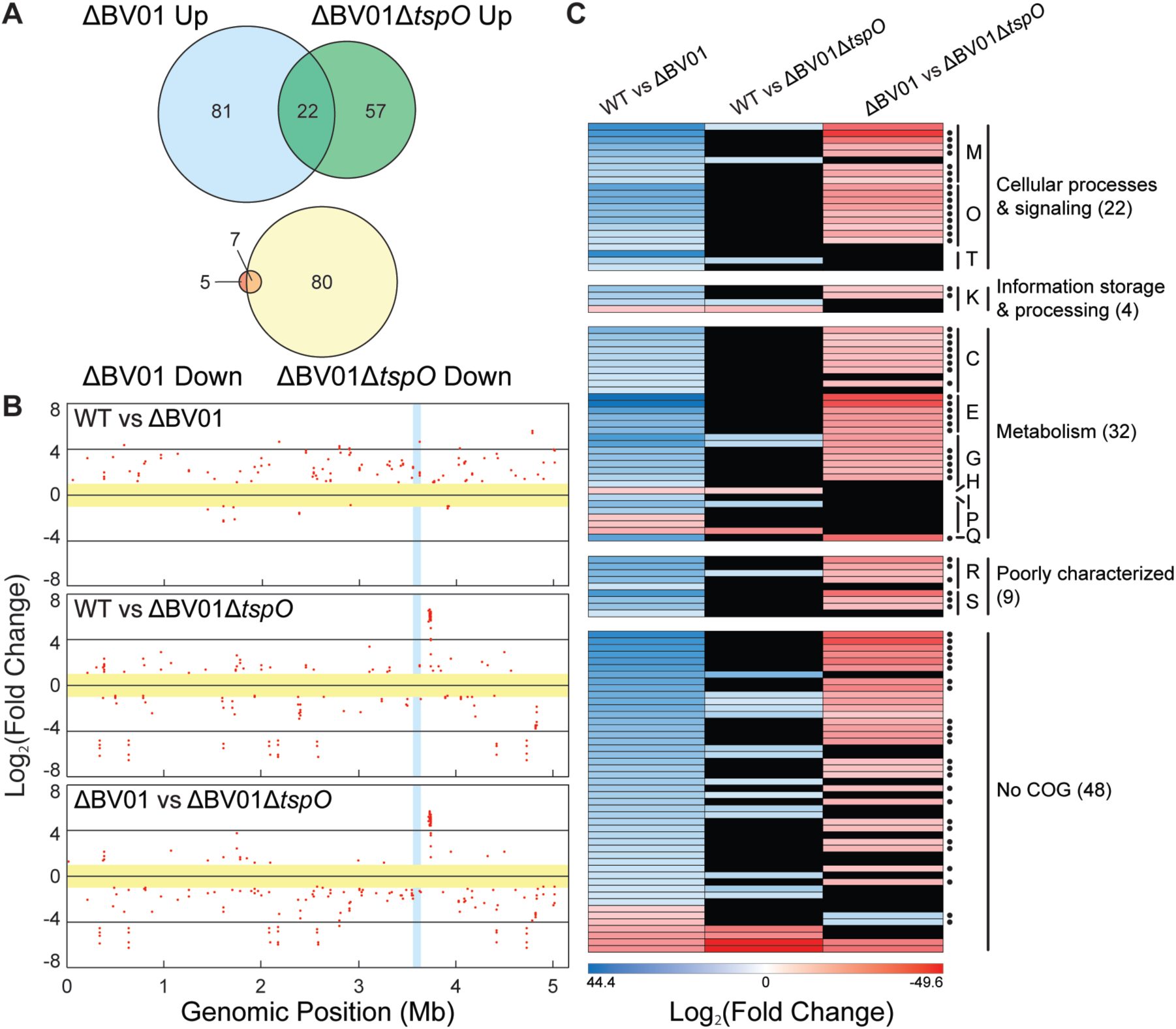
Differential regulation of the host transcriptome in response to BV01 lysogeny. (A) Count of differentially regulated transcripts as compared to the wild-type *B. vulgatus* lysogen (fold change ≥ 2, *q*-value ≤ 0.01). (B) Chromosomal localization of the differentially expressed genes. Each dot represents a differentially expressed transcript on a log_2_ scale; genes below the 2-fold change cutoff (yellow) and within the BV01 prophage (blue) not shown. Positive fold change values correspond to increased transcription in the second background listed. (C) General functional assignment of genes differentially expressed between wild-type and cured lysogen strains was accomplished using the Clusters of Orthologous Groups (COGs). Transcripts which are not differentially expressed in other strain comparisons are shown in black. *tspO*-dependent transcripts are marked on right (•). Positive fold change values correspond to increased transcription in the second background listed. Letters correspond to COG categories: cell wall/membrane/envelope biogenesis (M), post-translation modification, protein turnover, and chaperones (O), signal transduction mechanisms (T), transcription (K), energy production and conversion (C), amino acid transport and metabolism (E), carbohydrate transport and metabolism (G), coenzyme transport and metabolism (H), lipid transport and metabolism (I), inorganic ion transport and metabolism (P), secondary metabolite biosynthesis, transport, and catabolism (Q), general function prediction only (R), function unknown (S).

Analysis of RNAseq data revealed 115 host transcripts differentially regulated in response to lysogeny with BV01 (Fig. 3A), 103 of which (89%) are up-regulated in the cured lysogen (Table S1). These transcriptional changes occur across the host genome (Fig. 1B). Functional analysis of these transcripts revealed that most function in metabolism and cellular processes and signaling (Fig. 3C). Pathway analysis using the Kyoto Encyclopedia of Genes and Genomes Pathway Database (45), however, failed to yield pathway-level differences, which is likely a reflection of the level of annotation of the *B. vulgatus* genome. Taken together, these results indicate BV01 represses a diverse array of its host’s metabolic activities, suggesting it is acting through one or more transcriptional regulators.

One possible explanation for the widespread transcriptomic response to BV01 lysogeny is that a phage product directly alters the transcriptional activity of host genes. BV01 encodes two candidate genes that might act in this way: a predicted transcriptional regulator (BVU_RS14475) and a predicted sigma factor-like protein (BVU_RS14235) (Fig. S1). The transcriptional regulator encoded by BVU_RS14475 is the most highly transcribed gene in the BV01 prophage, encoding the putative CI-like repressor protein, which might directly interact with host promoters. Transcription of BVU_RS14235 is very low, so it is less likely to play a major role in the observed transcriptional response (Fig. S1). It is also possible that the observed transcriptional response to BV01 is the result of a host response to infection. A universal stress protein (*uspA*) homolog (BVU_RS16570) is upregulated in the BV01 lysogen, though it is not clear if that is a primary or secondary effect of infection.

### BV01 alters bile acid metabolism by disrupting the *tspO* promoter

Notably, integration of BV01 at the *attB* is correlated with a 23-fold down-regulation of the adjacent downstream transcript (BVU_RS14490), through an apparent disruption of the gene’s promoter (Fig. 4A). A low level of expression at this gene is observed in the wild-type lysogen, perhaps a result of readthrough from phage transcripts. This gene encodes a predicted tryptophan rich sensory protein (TspO) homolog (Fig. 4B), an intramembrane protein whose endogenous ligand is unknown, but which is broadly implicated in metabolic regulation and stress response in other bacteria (46–48). Although TspO is conserved in many bacteria, archaea, and eukaryotes, it is considered an accessory protein. Indeed, not all gut-associated members of the family Bacteroidales nor the genus *Bacteroides* encode *tspO* (Fig. 4C). Within the *Bacteroides*, *tspO* is restricted to the clade including *B. vulgatus* and *B. dorei*. Among *B. vulgatus* strains TspO is highly conserved (Fig. 4D), suggesting it plays a specialized role in regulating the cellular activities unique to *B. vulgatus*.

**Figure 4.**
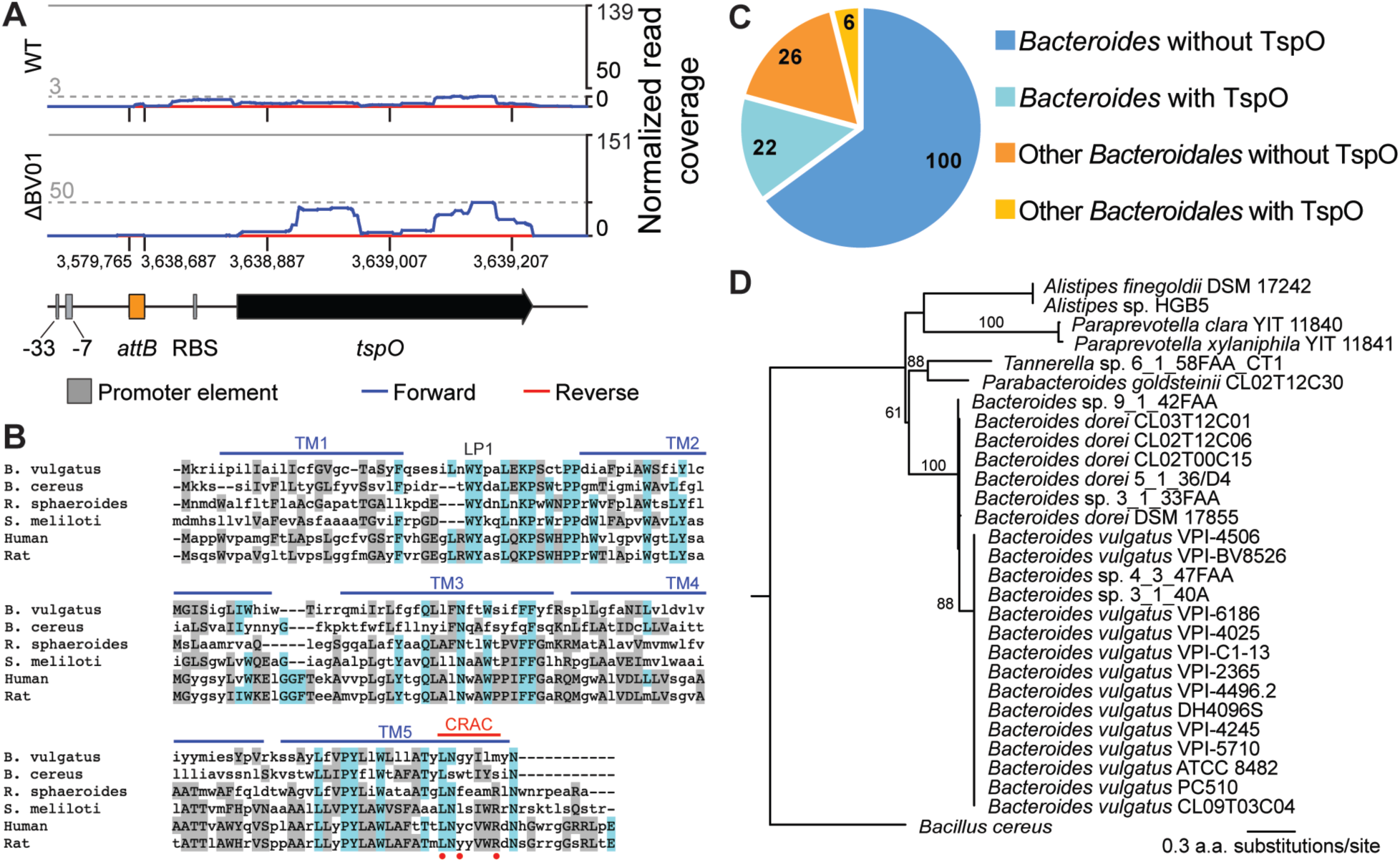
BV01 integration disrupts transcription of *tspO*. (A) Transcriptional activity of the *tspO* gene region as it exists in the cured lysogen background. RNAseq reads from wild-type (WT) and cured lysogen (ΔBV01) *B. vulgatus* were mapped to the region, and coverage was normalized to the total number of reads mapping to the genome. The average normalized read coverage for each genome is displayed as the y-axis maximum (grey line). Maximum read coverage for the region is displayed as the grey dashed line. (B) Amino acid alignment of *B. vulgatus* TspO with known TspO sequences was generated with MUSCLE (49). Identical and similar residues are colored blue and gray, respectively. Shown are TspO protein sequences from *B. vulgatus* (WP_005843416.1), *Bacillus cereus* (GCF80909.1), *Rhodobacter sphaeroides* (AAF24291.1), *Sinorhizobium meliloti* (AAF01195.1), human (NP_001243460.1), and rat (NP_036647.1). Secondary structures, cholesterol recognition/interaction amino acid consensus (CRAC) sequence, and critical residues (•) from *R. sphaeroides* TspO crystal structure are shown (50). (C) The search for TspO homologs in the family *Bacteroidales* was accomplished with a BLAST-based approach, using the *Bacillus cereus* copy of TspO (GCF80909.1) as a query against a database of 154 gut-associated *Bacteroidales* genomes, 122 of which are from the genus *Bacteroides*. Genome counts are indicated within categories. (D) Gene tree estimated from TspO sequences across the *Bacteroidales*. All *B. vulgatus* and *B. dorei* genomes included in the search encode *tspO*. Clade for *B. vulgatus* TspO sequences is displayed as a polytomy; all *B. vulgatus* TspO sequences are at least 98% identical to each other. Numbers above branches represent bootstrap values; only bootstraps over 50 shown. The gene tree was estimated using FastTree (51).

Given TspO’s important role in regulating cellular activities in other bacterial systems, it was hypothesized that it may be responsible for some of the differential regulation observed in response to prophage BV01. A *tspO* deletion mutant was constructed in the cured lysogen background (ΔBV01Δ*tspO*) and its transcriptome sequenced alongside that of the wild-type and cured lysogen strains (Fig. 3). The predicted *tspO* regulon extends far beyond the differential expression observed in the cured lysogen (Fig. 3A, 3B), suggesting the small amount of *tspO* transcription in the BV01 lysogen exerts effects on the rest of the genome. Transcripts differentially regulated between the BV01 wild-type lysogen and cured lysogen which returned to wild type-like levels upon further deletion of *tspO* were classified as *tspO*-dependent transcripts (Fig. 3C). Of the 115 transcripts differentially regulated in response to BV01, 69 (60%) are *tspO*-dependent. Consistent with TspO’s role in regulating stress, many *tspO*-dependent transcripts fall into the COG category for post-translation modification, protein turnover, and chaperones, including several thioredoxins, peroxidases, and protein chaperones. *tspO*-dependent transcripts also account for the majority of metabolic genes differentially regulated in response to BV01.

Two *tspO*-dependent transcripts that are down-regulated in response to lysogeny with BV01 encode putative bile salt hydrolases (BVU_RS13575, 5.38-fold change, *q* < 10^-100^; BVU_RS20010, 6.58-fold change, *q* < 10^-200^). It was hypothesized these transcriptional differences might reflect enzyme activities. To this end, *B. vulgatus* strains were grown in the presence of bile acids and the deconjugation of those bile acids measured by LC/MS (Fig. 5).

**Figure 5.**
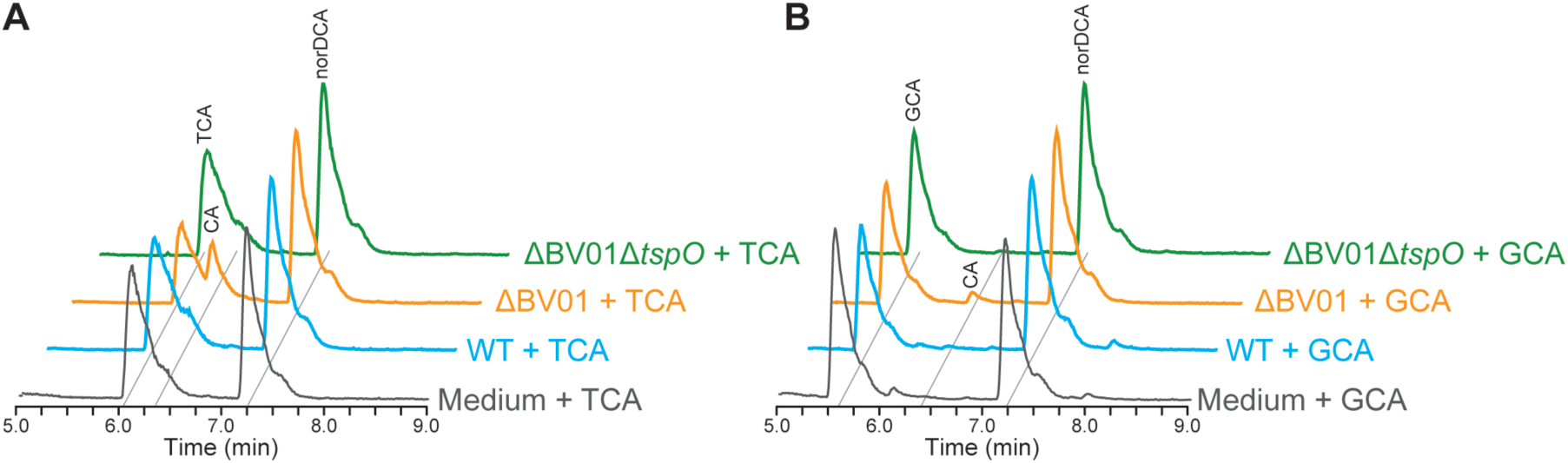
BV01 alters host interactions with bile acids in a *tspO*-dependent manner. (A) Representative LC/MS traces showing *B. vulgatus* deconjugates taurocholic acid (TCA) to cholic acid (CA) in the cured lysogen background (ΔBV01), but little or no activity CA is detectable in the wild-type (WT) or cured lysogen *tspO* deletion (ΔBV01Δ*tspO*) backgrounds. *B. vulgatus* cultures were incubated with 50 μM TCA for 16 hr prior to bile acid extraction. (B) Representative LC/MS traces showing *B. vulgatus* deconjugates glycocholic acid (GCA) to CA in the ΔBV01 background, but not in the WT or ΔBV01Δ*tspO* backgrounds. *B. vulgatus* cultures were incubated with 50 μM GCA for 48 hr prior to bile acid extraction. Nordeoxycholic acid (norDCA) was added to a final concentration of 15 μM as an internal standard after incubation. Peaks labeled for their metabolites based on m/z; TCA = 514.29, GCA = 464.30, CA = 407.28, norDCA = 377.27.

LC/MS results show that the wild-type *B. vulgatus* lysogen does not significantly deconjugate glycocholic acid (GCA) to cholic acid (CA), and may exhibit modest deconjugation of taurocholic acid (TCA) (Fig. 5). This agrees with a previous study that showed *B. vulgatus* ATCC 8482 can deconjugate TCA but cannot deconjugate GCA over a 48 hr incubation (49). Importantly, CA is clearly detectable only in the cured lysogen background, consistent with our prediction based on RNAseq data. This bile acid deconjugation phenotype is ablated with the deletion of *tspO*, further supporting the hypothesis that *tspO* activates transcription of bile salt hydrolases, resulting in their increased enzymatic activity.

To see if *tspO*-disrupted lysogens occur in natural human gut microbiomes, read mapping from 256 healthy human gut metagenomes was performed. Starting with reads which mapped to *tspO* in the reverse orientation, read mates were checked for mapping to Phage BV01 and its relatives (Fig. 6). All samples had reads mapping to *tspO*, with an average of 0.0004% of metagenome reads mapping, indicating the corresponding population of *B. vulgatus* and *B. dorei* encoding *tspO* is relatively abundant (Fig. S2A). Incidence of *tspO* associated with BV01 or a related phage is also common; 13.3% of samples (*n*=34) contained read pairs mapping to *tspO* and an adjacent prophage. Within an individual microbiome, incidence of *tspO*-disrupted lysogens appears rare, usually comprising 3% or less of the combined *B. vulgatus* and *B. dorei* population, although for some individuals this incidence rate can be more than 10% (Fig. S2B). In addition to these patterns, carriage of *tspO*-disrupted lysogens appears stable over time, as indicated by individuals sampled at multiple time points (Table S2). Together, these data suggest that BV01 and other phages’ effects on downstream phenotypes via *tspO* are likely quite common among humans, though these lysogens comprise the minority of the overall microbiome.

**Figure 6.**
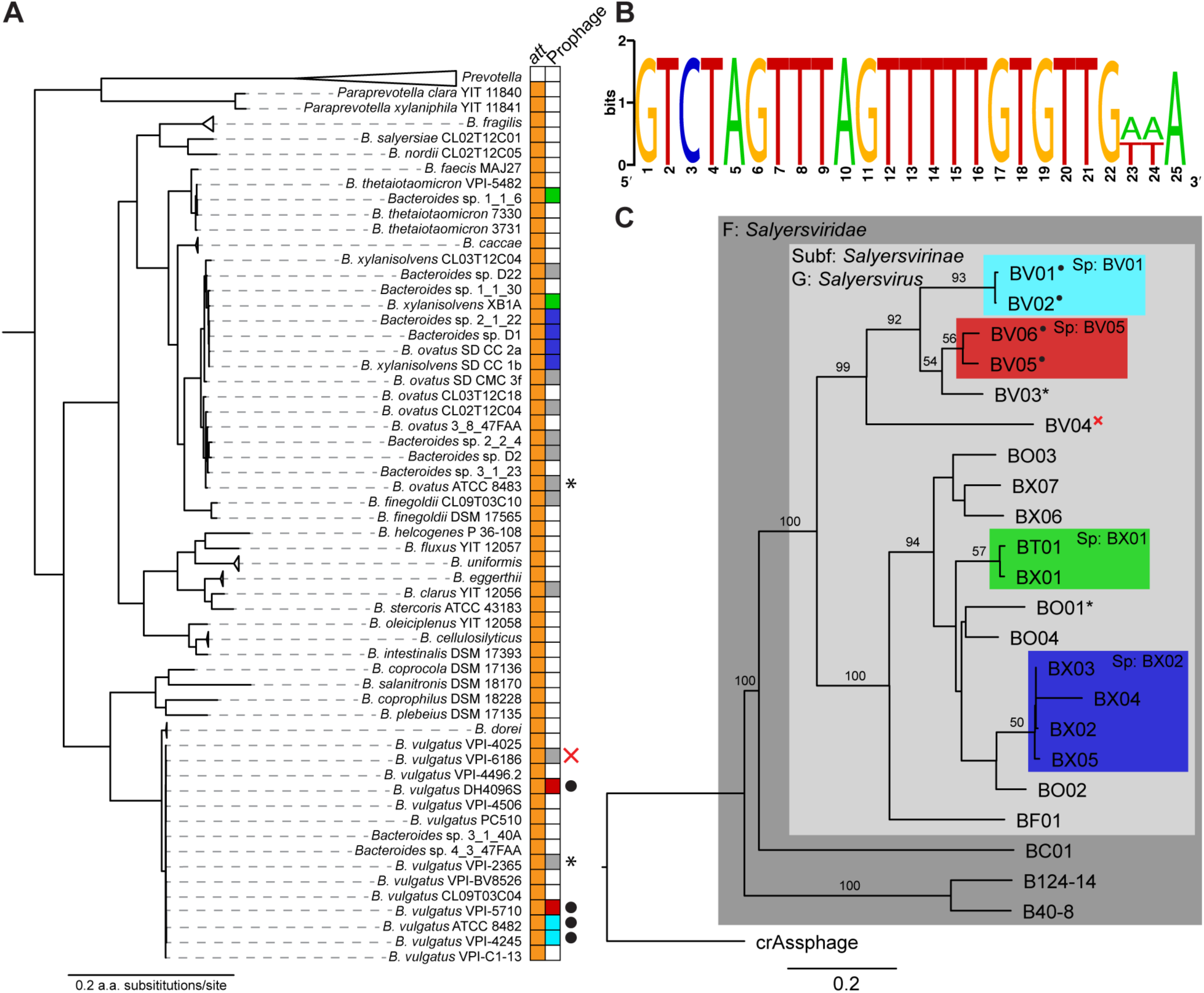
*Salyersviridae* occur throughout the *Bacteroides* genus. (A) *Bacteroides* phylogeny and occurrence of *Salyersviridae att* site. All duplications of the *att* site are associated with a putative integrated prophage. Host phylogeny estimated by maximum likelihood from concatenated alignment of 13 core genes. (B) Consensus *att* site for *Salyersvirinae*. The *attP* is duplicated upon integration of a *Salyersvirinae* prophage, resulting in direct repeats. Image made with the WebLogo online tool. (C) Phylogenomic Genome-BLAST Distance Phylogeny implemented with the VCTOR online tool (53) using amino acid data from all phage ORFs. For consistency, all phage genomes were annotated with MetaGeneAnnotator (54) implemented via VirSorter (55). Support values above branches are GBDP pseudo-bootstrap values from 100 replications. Family (F), subfamily (Subf), genus (G), and species (Sp) assigned by OPTSIL clustering (56) (Table 1). Each leaf of the tree represents a unique phage species, except where indicated by colored boxes. Active prophages confirmed by sequencing and/or PCR indicated with “•”; prophages confirmed to have been inactivated by genome rearrangement indicated with “x”; prophages which were tested for activity with inconclusive results indicated with “*” (Fig. S4).

**Table 1.**
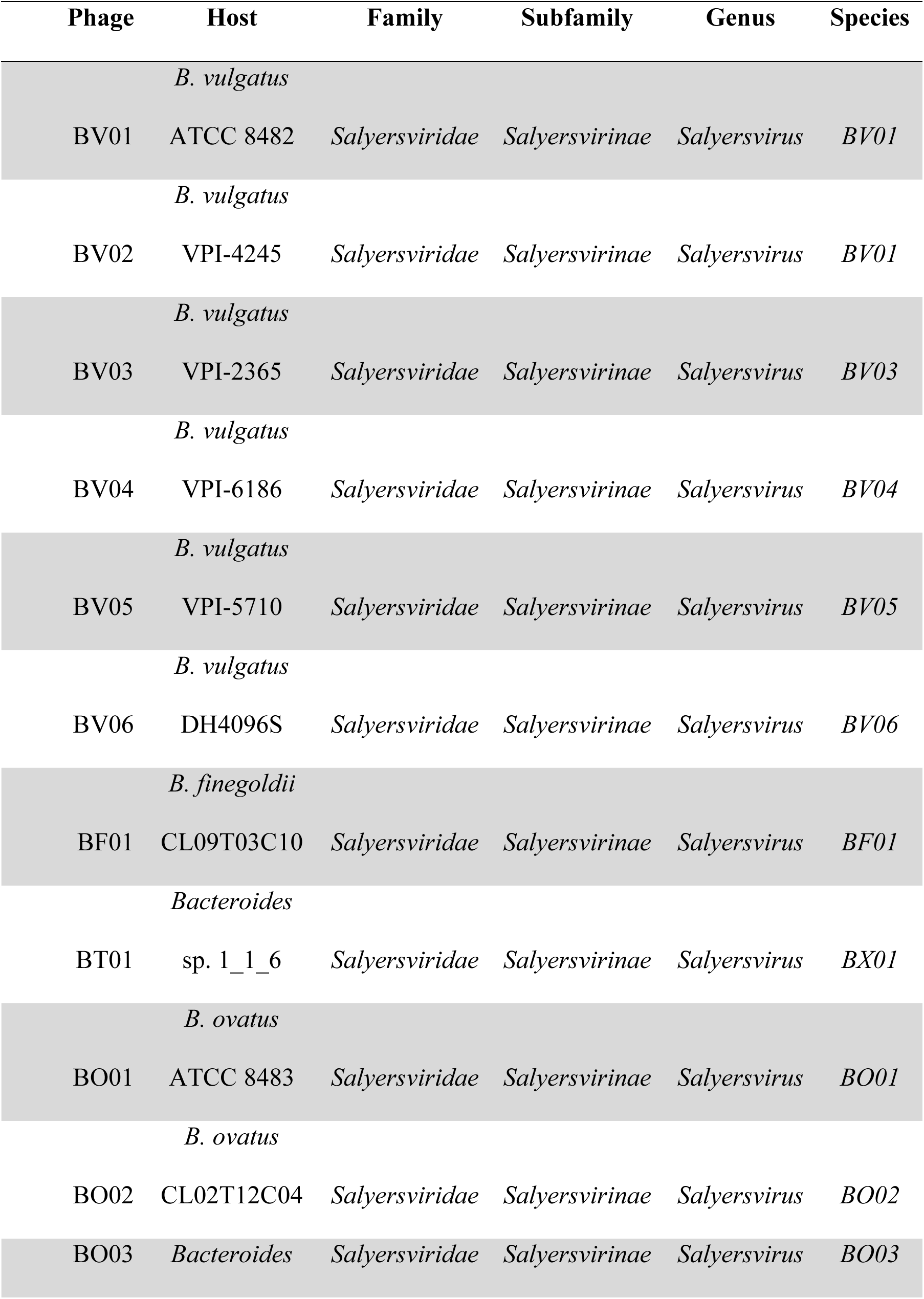
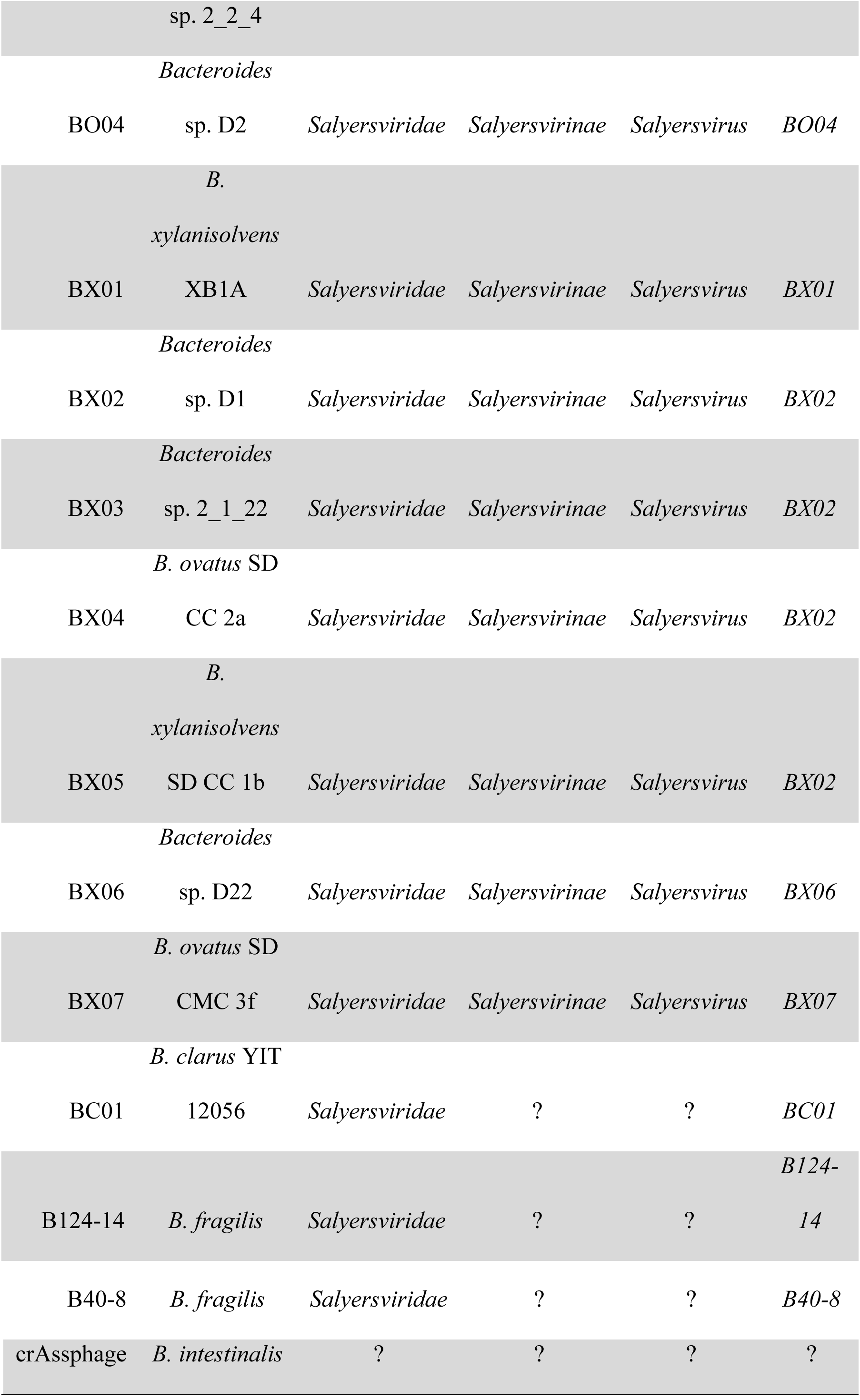
Bacteroides phage taxonomy as determined by whole genome clustering.

### BV01 represents the proposed viral family *Salyersviridae*

While searching for potential new hosts for BV01, its predicted 25-bp *attB* was queried against 154 gut-associated Bacteroidales genomes. It was found that all *Paraprevotella* and *Bacteroides* genomes had at least the first 21 bases of the attachment site conserved (Fig. 6A). In 20 genomes, two copies of the *att* site were found, and all are associated with putative prophages. In all instances, the putative *attL* and *attR* sites are direct repeats, as is true for BV01. Importantly, only *B. vulgatus* lysogens encode *tspO* (Fig. 4); other prophages and *attB* sites occur in an alternative genomic context. Alignment of these putative prophage-associated *att* sites finds that the first 22 basepairs are always conserved, and an additional 3 basepairs are variable among lysogen genomes (Fig. 6B).

Integration at the same *att* site suggests these prophages are genetically related. To assess relatedness, VICTOR (50) was used to build a genome tree from all of the identified prophages (Fig. 6C) and OPTSIL clustering (51) was used to predict taxonomic groups, which we named *Salyersviridae*, *Salyersvirinae*, and *Salyersvirus* (Table 1). Taxonomic clustering also defined phage species, four of which have more than one member. Interestingly, the phage species BX01 has representatives in two distantly related hosts, *Bacteroides xylanisolvens* XB1A (phage BX01) and *B. thetaiotaomicron* 1_1_6 (phage BT01), suggesting that at least some Salyersviruses have broad host ranges. Phage BC01 was placed outside the subfamily *Salyersvirinae*, which is consistent with its considerable sequence divergence (Fig. S3). Further examination of the BC01 *attL* and *attR* sites found that the 22-bp consensus sequence is only a portion of the full 69-bp repeat flanking the BC01 prophage, which further supports placement of BC01 outside the *Salyersvirinae*. Also included in the analysis were three outgroup sequences from the known *Bacteroides*-infecting lytic phages: B124-14, B40-8, and crAssphage. Surprisingly, OPTSIL taxonomic clustering placed phages B124-14 and B40-8 in the family *Salyersviridae*. This similarity is not detectable at the level of nucleotide sequence (Fig. S3), but is likely driven by similarities in several proteins, including homologs to the predicted lysin, excisionase, and ssDNA binding protein from BV01.

To check that the proposed *Salyersviridae* clade was comprised of active phages, we searched for evidence of activity in culture and in wastewater samples. Sequencing and PCR were used to test the activity of six additional Salyersviruses (Fig. S4). Phages BV02, BV05, and BV06 were found to be released from host cells in culture, while BV03 is likely inactivated by genomic rearrangement. We did not observe activity for BV04 or BO03 from *B. ovatus* ATCC 8483, both of which appear complete based on synteny with intact Salyersviruses. This may be the result of inactivating mutations or not being induced in the growth conditions tested (Fig. S4D).

Furthermore, we used read mapping to search for evidence of all 20 *Salyersviridae* phages in a wastewater metavirome (Fig. S5A). Reads mapped to all *Salyersviridae* genomes, however it may be the result of sequence conservation within the family, as reads often accumulate at the most conserved regions of each phage genome. Further, very little read mapping occurred in the distal portion of the inactivated prophage BV04, which resembles host chromosome more than phage sequence, indicating that virome processing removed most cellular DNA prior to sequencing (Fig. S5A). *De novo* assembly of the individual wastewater viromes finds contigs which align with high identity but imperfectly to each *Salyersviridae* phage, supporting the findings seen by read mapping, and suggesting the real diversity of the family *Salyersviridae* is far greater than what has been observed integrated in cultured host genomes so far. This analysis found that phages infecting *B. vulgatus* are more abundant than other *Salyersviridae* phages based on maximum normalized read coverages. A similar comparison concludes that most individual temperate *Salyersviridae* phages are approximately equal in abundance to the lytic *Salyersviridae* phage B124-14 and B40-8, and approximately ten-fold less abundant than crAssphage, in these wastewater viromes (Fig. S5A). Searches for *Salyersviridae* phages in individual healthy human fecal metagenomes refine this conclusion, showing that in most human samples, *Salyersviridae* phages are at least as abundant as crAssphage, though crAssphage can reach very high abundances in a subset of individuals (Fig. S5B). Although confirmation of most *Salyersviridae* activities will require better sampling and *in vitro* testing, these results indicate the phage family is active in human-associated communities.

## DISCUSSION

Here, we characterize a complex phage-host interaction between *Bacteroides phage BV01* and its host *B. vulgatus.* We first demonstrate that BV01 is an intact prophage capable of directing its own excision, and it is transducible *in vivo* in a gnotobiotic mouse model. Using a combination of genetics, RNAseq, and analytical chemistry, we show that BV01 decreases its host’s ability to deconjugate bile acids by disrupting the transcription of the gene adjacent to the *attB* encoding a TspO homolog. Furthermore, we show that *tspO* disruption by phage integration is common among, but rare within, healthy human gut microbiomes, and can be mediated by BV01 or its relatives. Together, these findings elucidate a complex mechanism by which a phage alters its host’s activities.

The repression of bile acid deconjugation as a consequence of BV01 integration is particularly relevant in the context of the mammalian gut. Mammals secrete conjugated primary bile acids into the small intestine, where they reach concentrations as high as 1 mM (52); though the majority of bile acids secreted into the small intestine are readsorbed, they can still accumulate to concentrations of 0.2-1 mM in the colon (53). While bile acids are broadly capable of damaging lipid membranes, generally *Bacteroides* species are considered bile-resistant (54), the mechanism of which is unknown. Bile acid deconjugation is a common activity encoded by gut-associated microbes, though its direct benefit to those microbes is unclear. Microbial modification of the bile acid pool can be linked to beneficial changes in the human host metabolism (49, 55) and varied epithelial susceptibility to viral pathogens (56). The link between BV01 and bile acid metabolism suggests a heretofore undescribed mechanism by which gut phages might influence mammalian host phenotypes.

Here, bile acid deconjugation in *B. vulgatus* is dependent on a putative TspO. Bacterial TspOs are important for regulating metabolic switches and stress regulation in at least three diverse systems (46–48), though the mechanism of action for the protein is unknown. The crystal structure of TspO shows a periplasm-facing binding pocket distinct from the intramembrane cholesterol recognition consensus sequence, which may bind or degrade porphyrins (57–59). Both the porphyrin degrading and cholesterol transporting functions of TspO, however, have been disputed (60, 61). Despite this, it is notable that cholesterol is structurally similar to bile acids, being their biosynthetic precursor. In at least one other gut-associated microbe, *tspO* is up-regulated by bile acids, suggesting TspO may be involved in bile acid metabolism in gut microbes more broadly (62). The regulatory link described here between bile acid hydrolysis and TspO suggests a hypothesis where the *B. vulgatus* TspO might be a sensor and regulator of bile acid interactions.

Induction of BV01 from its integrated state and infection of new hosts remains enigmatic. Prophage induction is canonically linked to stress-dependent pathways, as is the case for lambdoid phages that respond to DNA damage via RecA-dependent cleavage of the CI repressor protein (63). It is possible that prophages in *Bacteroides* hosts respond to alternative stimuli, as is the case for CTnDOT, a well-studied *Bacteroides* conjugative transposon, whose excision is inducible only by teteracycline (64). Neither DNA damage nor antibiotics induce prophage BV01 *in vitro*, so all experiments here relied on an apparently low rate of spontaneous prophage induction. Similarly, no infecting conditions or susceptible hosts have been identified for BV01 *in vitro.* We demonstrate that BV01 is transducible in a gnotobiotic mouse model, suggesting that an unknown mammalian host factor is required for novel BV01 infection. Enigmatic infection dynamics may be the result of the phase variable polysaccharide capsule, as recent work suggests heterogeneity in capsular composition hinders phage infection on population-scales (17). Indeed, it has long been observed that finding phages in the *Bacteroides* using traditional techniques is difficult or impossible for most host strains (65, 66), making the host-first approach to phage discovery used here especially appealing.

Finally, phage BV01 is the first representative of a broad family of phages that spans an entire host genus, and includes lytic and temperate members. *Salyersviridae* is common and diverse among natural human samples, but rare within individuals, suggesting lysogenization may confer frequency-dependent advantages to the bacterial host. The genetic context of non-*B. vulgatus Salyersviridae* lysogens remains unexplored, providing ample opportunity for further discovery of novel phage-host interactions. The absence of *tspO* in these other host systems may provide the ideal background for studying more direct impacts of these phages on their hosts. Certainly, other interactions between BV01 and its host remain to be studied, though they were overshadowed here by the enormous effects of *tspO*. Future studies should also examine the role of *Salyersviridae* phages on bacterial host fitness and evolution (67, 68), as these phages directly impact their bacterial hosts and those interactions likely have important ripple effects throughout the microbiome and on the mammalian host that remain to be elucidated.

## MATERIALS AND METHODS

### Strains and culture conditions

All strains and plasmids used in the study are listed in Table S3. *Escherichia coli* S17-1 *λ pir* was used for all routine recombinant DNA cloning, and grown aerobically in Lysogeny Broth (LB) at 37°C. *B. vulgatus* strains were cultured anaerobically in a vinyl anaerobic chamber using 70% N_2_, 20% CO_2_, and 10% H_2_ gas mixture (Coy Laboratory Products, Grass Lake, MI). All *B. vulgatus* cultures were grown on Difco Brain Heart Infusion (BHI) agar supplemented with 10% defibrinated horse blood (BHI-HB; Quad Five, Ryegate, MT), or in tryptone-yeast extract-glucose (TYG) broth (69) at 37°C. When necessary, ampicillin (100 µg/mL), gentamicin (200 µg/mL), erythromycin (25 µg/mL), 5’-flourodeoxyuridine (FUdR; 20 µg/mL), or tetracycline (2 µg/mL) were supplemented in the media. Infection assays were performed on BHI supplemented with 50 μg/mL hemin and 0.5 μg/mL menadione (BHI-HM) and TYG_S_ (70).

### Genetic manipulation

All primers used to construct genetic mutants are listed in Table S3. Markerless deletion mutants in *B. vulgatus* were achieved by allelic exchange using a system analogous to that developed in *B. thetaiotaomicron,* (71) and confirmed by PCR and whole genome sequencing. The *tdk* gene (BVU_RS09305), encoding thymidine kinase, was deleted from *B. vulgatus* ATCC 8482 by allelic exchange, conferring resistance to the toxic nucleotide analog FUdR. Cloning was performed as described by Degnan *et al.* (72). Briefly, the 3.5 Kb regions flanking either side of *tdk* were amplified with Kapa HiFi Taq MasterMix (Kapa Biosystems, Wilmington, MA) and joined by splicing overlap exchange (SOE) PCR. The SOE product was purified, restriction digested and ligated into the suicide vector pKNOCK-*bla*-*ermG*b in *E. coli*, and conjugated into *B. vulgatus.* Single recombinant merodiploids were selected for on BHI-HB supplemented with gentamicin and erythromycin, and double recombinant deletion mutants subsequently selected for on BHI-HB with FUdR. The counterselectable suicide vector pExchange-*tdk*BV was constructed by amplifying *tdk* from *B. vulgatus* and cloning the *tdk* amplicon into pKNOCK-bla-ermG_b_ by the same methods used to clone the SOE product above.

Subsequent deletions were accomplished similarly as described for *tdk*, except using pExchange-*tdk*BV and flanking regions of ∼1 Kb to create the deletion alleles (*tspO, int*). For deletion of the entire BV01 provirus, an empty attachment site (*attB*) and the flanking 800 bp were cloned from *B. vulgatus* VPI-4506, which has 99.9% nucleotide identity to the analogous regions flanking BV01 in *B. vulgatus* ATCC 8482.

Complementation of the BV01 integrase (BVU_RS14130) was accomplished by cloning the gene and its native promoter into the integrative plasmid pNBU2-*bla*-*ermG*b, which has a single integration site in the *B. vulgatus* genome (*attN*; NC_009614.1:3152550..3152572). This construct was conjugated into *B. vulgatus* and transconjugants selected for on BHI-HB supplemented with gentamicin and erythromycin as described elsewhere (72).

The BV01-*tetQ* strain was constructed by inserting *tetQ* from pNBU2-*bla*-*tetQ* immediately downstream of the stop codon of BVU_RS14265, upstream of a predicted transcriptional terminator. As was done for deletion constructs, the desired region was constructed on the pExchange-*tdk*BV plasmid and moved into the wild-type *B. vulgatus* strain by allelic exchange. First, a ∼2 Kb region surrounding the BVU_RS14265 stop codon was amplified in two pieces with SOE primers designed to insert adjacent SpeI and BamHI cut sites downstream of the stop codon and ligated into pExchange-*tdk*BV. This construct was confirmed by Sanger sequencing before *tetQ* and its promoter were amplified from CTnDOT, and ligated into the SpeI and BamHI cut sites. Tetracycline was used to select for mutants, and release of BV01-*tetQ* phages confirmed by PCR.

Select mutant strains were confirmed by whole genome sequencing and analyzed with Breseq (73) aligned to the wild-type *B. vulgatus* ATCC 8482 genome (NC_009614.1), and summarized in Table S4.

### Genome sequencing

Cells were pelleted from 5 mL overnight culture in TYG by centrifugation at 4,000 × *g* for 5 min at 4°C, resuspended in 0.5 mL TE buffer (10 mM Tris, 1 mM EDTA), and lysed by adding sodium dodecyl sulfate (SDS) and proteinase K (GoldBio, Olivette, MO) to final concentrations of 0.07% and 300 µg/mL, respectively, and incubating for 2 hr at 55°C. Cellular material was removed by washing twice in an equal volume of buffered phenol, phenol-chloroform-isoamyl alcohol (VWR, Radnor, PA), and DNA precipitated with 100% ethanol in the presence of 0.3M sodium acetate at −20°C overnight. DNA pellets were washed with 70% ethanol, dried, and resuspended in TE buffer.

Phage DNA was prepared from overnight TYG culture supernatants collected after centrifugation and concentrated by centrifugation with 30,000 MWCO Corning Spin-X UF 20 Concentrators (Corning, NY) or by tangential flow filtration with a Vivaflow 50R 30,000 MWCO Hydrosart filter (Sartorius, Gottingen, Germany). Supernatants were treated with 200 µg/mL DNase I and 1 µg/mL RNase A for 1 hr at room temperature to remove unprotected DNA and RNA. Virions were disrupted with 1% SDS and 1 mg/mL proteinase K for 2 hr at 55°C. DNA was further isolated using the same phenol-chloroform-isoamyl alcohol extraction and ethanol precipitation procedures as for cellular DNA.

DNA libraries were constructed with the Nextera XT Library Preparation Kit and Index Kit (Illumina, San Diego, CA). DNA libraries were pooled and sequenced on both the Illumina HiSeq5000 and HiSeq2500 and fastq files were generated from demultiplexed reads with bcl2fastq Conversion Software (Illumina, San Diego, CA). Reads were trimmed and assembled using the A5ud pipeline (74). Sequencing methods and assembly data are summarized in Table S5.

### Genome annotation

Annotation of cellular genomes was accomplished with a custom Perl script that calls protein coding genes with Prodigal (75) and RNA coding genes with tRNAscan-SE (77), Rnammer (78) and Infernal (78). Functional predictions are assigned by searching against Kyoto Encyclopedia of Genes and Genomes (45), Cluster of Orthologous Genes (80), Pfam (80), and TIGRFAM (81) databases, and subCELlular LOcalization predictor (82) is used to predict cellular localization.

For phage genomes, genes were called by Prodigal and the gene calling tool within Artemis (83). Functional predictions were made as above except with relaxed search parameters (cut_tc in hmmscan), plus using Basic Local Search Alignment Tool (84) with the Genbank virus database (85), Phyre2 (86) to identify conserved protein folds, and iVireons (87) to predict structural proteins, and manually comparing and combining results.

### Integrase activity assays

Integrase activity was assayed through PCR of DNase-treated supernatant DNA. Briefly, free phage DNA was prepared as for DNA sequencing, and amplified with Kapa HiFi Taq MasterMix with primers specific to BV01 (BVU_RS14350) or spanning the circularized *attP* (Table S3). Free phage DNAs were checked for the presence of contaminating cellular DNA by amplifying the 16S rRNA gene with universal primers. Amplicons were cleaned with a Qiagen PCR Cleanup kit (Hilden, Germany) and run on an agarose gel in 0.5X Tris-borate-EDTA buffer at 70V alongside 1 Kb ladder (New England BioLabs, Ipswich, MA) or GeneRule Express DNA Ladder (Thermo Scientific, Waltham, MA) and stained with GelRed (VWR, Radnor, PA). Amplicons generated with *attP*-flanking primers were sequence confirmed by Sanger sequencing performed by ACGT, Inc (Wheeling, IL).

### Gnotobiotic mice

All experiments using mice were performed using protocols approved by the University of California Riverside Institutional Animal Care and Use Committee. Germfree C57BL/6J mice were maintained in flexible plastic gnotobiotic isolators with a 12-hr light/dark cycle. Animals caged individually (*n*=1, female) or in pairs of litter mates (*n*=6, males) were provided with standard, autoclaved mouse chow (5K67 LabDiet, Purina) *ad libitum*. With no *a priori* reason to expect age to influence transduction rates, animals ranged from 7 weeks to nearly 12 months old. Individually antibiotic resistance marked bacterial strains were grown individually for ∼20h in TYG medium with appropriate antibiotics and frozen at −80°C in anaerobic cryovials. Cell viability was tested by plating and viable CFU counts were used to combine equal parts of the wild-type *B. vulgatus* lysogen tagged with pNBU2-*bla*-*ermGb* and *B. vulgatus* BV01-*tetQ* tagged with pNBU2-*bla*-*cfx*. Approximately 4 x 10^6^ CFUs of the combined strains were administered to each animal by oral gavage. Fecal samples were collected on days 1, 3, 7 and 11 from each animal. Fecal pellets were processed by adding 500µl of TYG+20% Glycerol to each tube and vigorously shaking in a beadbeater without beads for 1m 30s. Fecal slurries were spun down at 2,000 x g for 1 s, followed by serial dilution on selective media (BHI+Tet+Gn, BHI+Erm+Gn, BHI+Erm+Tet+Gn) to determine CFUs. Animals were sacrificed on d11 following final fecal collection.

### Transcriptomic response to lysogeny

*B. vulgatus* was grown overnight in 5 mL TYG medium. Each culture was pelleted (4,000 × *g* for 5 min at 4°C), supernatant decanted, and washed in an equal volume of TYG three times. Cells were normalized to an OD_600_ of ∼0.3 and used to inoculate cultures in 10 mL TYG at a final dilution of 1:1000 in biological triplicate. Cell growth was monitored and cells were harvested at an OD_600_ of ∼0.4. Total RNA was prepared with a Qiagen RNeasy kit (Hilden, Germany) and treated on-column with RNase-free DNase (Qiagen, Hilden, Germany). RNA was quantitated with a Qubit 2.0 fluorometer (Thermo Fisher, Waltham, MA) and stored at −80°C.

RNA was submitted to the W. M. Keck Center for Comparative and Functional Genomics at the University of Illinois at Urbana-Champaign for quality analysis, rRNA depletion with the RiboZero Bacteria kit (Illumina, San Diego, CA), library construction with the TruSeq Stranded mRNAseq Sample Prep kit (Illumina, San Diego, CA), and sequencing on an Illumina NovaSeq 6000 with the NovaSeq S4 reagent kit. Fastq files of demultiplexed reads were prepared with the bcl2fastq v2.20 Conversion Software (Illumina, San Diego, CA).

RNAseq reads were quality filtered and trimmed with trim_galore v0.4.4 (https://www.bioinformatics.babraham.ac.uk/projects/trim_galore/). Rockhopper (88) was used to identify differentially expressed transcripts between isogenic mutants (≥2-fold change, *q* ≤ 0.01).

### Bile salt deconjugation assay & LC/MS

*B. vulgatus* strains were inoculated in TYG liquid supplemented with 50 μM glycocholic acid (GCA) or 50 μM taurocholic acid (TCA) and allowed to grow for 16 or 28 hr, respectively. Grown cultures were brought to a pH 2.0-3.0 with 10 N hydrochloric acid, centrifuged for 5 min at 4,000 x *g*, and the pellets discarded. Bile acids were isolated by solid phase extraction over Sep-Pak tC18 500 mg cartridges (Waters Corp., Milford, MA). Cartridges were preconditioned by serial washes with 6 mL hexane, 3 mL acetone, 6 mL methanol, and 6 mL water (pH = 3.0). Acidified supernatants were loaded before washing with 3 mL 40% methanol. The column was allowed to dry, then bile acids eluted in 3 mL methanol. Samples were evaporated under nitrogen, resuspended in 20 μL methanol, and centrifuged before analysis by liquid chromatography-mass spectroscopy (LC/MS).

LC/MS for all samples was performed on a Waters Aquity UPLC coupled with a Waters Synapt G2-Si ESI MS. Chromatography was performed using a Waters Cortecs UPLC C18 column (1.6 µm particle size) (2.5 mm x 50 mm) with a column temperature of 40° C. Samples were injected at 1 µl. Solvent A consisted of 95% water, 5% acetonitrile, and 0.1% formic acid. Solvent B consisted of 95% acetonitrile, 5% water, and 0.1% formic acid. The initial mobile phase was 90% Solvent A, 10% Solvent B and increased linearly until the gradient reached 50% Solvent A and 50% Solvent B at 7.5 min. Solvent B was increased linearly again until it was briefly 100% at 8.0 min until returning to the initial mobile phase (90% Solvent A, 10% Solvent B) over the next 2 min. The total run was 10 min with a flow rate of 10 µL/min. MS was performed in negative ion mode. Nebulizer gas pressure was maintained at 400° C and gas flow was 800 L/hour. The capillary voltage was set at 2,000 V in negative mode. MassLynx was used to analyze chromatographs and mass spectrometry data.

### Taxonomic nomenclature

The family, subfamily, and generic names were chosen to honor the microbiologist Abigail A. Salyers, who made significant contributions to the understanding of function and genetics of human gut anaerobes and the importance of their mobile genetic elements.

### Wastewater collection, processing, and viromics

From the Urbana & Champaign Sanitary District Northeast Plant (Urbana, IL), 1 L of unprocessed wastewater was collected at each of three time points: May 25, 2016, June 23, 2016, and October 3, 2016.

Wastewater samples were transported on ice, immediately centrifuged at 2,500 ×*g* for 10 min at 4°C and filtered through a 0.4 μm polyethersulfone filter to remove large particulate and cellular matter. The sample was split into three aliquots and processed three ways. One aliquot was not processed further (F). Another aliquot was filtered a second time through a 0.22 μm polyethersulfone filter (DF). The last aliquot was washed three times with an equal volume of chloroform (FC). All aliquots were concentrated 100-fold and virome DNA was isolated from each as described for genome sequencing of phages.

DNA libraries of virome DNA were prepared using the same methods as described for genome sequencing and were sequenced on an Illumina HiSeq 2500 sequencer with a HiSeq v4 SBS sequencing kit (Illumina, San Diego, CA) producing 2×160-bp paired-end reads. Fastq files of demultiplexed reads were generated with the bcl2fastq v2.17.1.14 Conversion Sotfware (Illumina, San Diego, CA). Reads were trimmed and quality filtered using Trimmomatic 0.38 (89) and assembled with metaSPAdes v3.13.0 using default parameters (90). Sequencing and assembly data for wastewater viromes is summarized in Table S5. Read mapping to phage genomes was performed with bwa (91).

Human Microbiome Project Healthy Human Subjects Study samples were downloaded with portal_client (Table S2). Read mapping was performed with bwa (91).

### Data availability

Trimmed RNAseq reads from this study are deposited in the NCBI SRA under PRJNA622597; sample accession numbers are SAMN14522273 for the wild-type lysogen (WT), SAMN14522274 for the cured lysogen (ΔBV01), and SAMN14522275 for the cured lysogen *tspO* deletion (ΔBV01Δ*tspO*). Trimmed wastewater virome reads are deposited in the NCBI SRA under PRJNA622299. Ten new assembled *B. vulgatus* genomes are deposited in NCBI GenBank under PRJNA622758.

## ACKNOWLEDGMENTS

We thank Nadja Shoemaker and Abigail Salyers for access to an impressive collection of *Bacteroides* isolates; Ken Ringwald for critical review of the manuscript; Jim Imlay for insightful discussion on metabolism and stress; Alvaro Hernandez and Chris Wright for DNA and RNA sequencing; Bruce Rabe for aid in wastewater collection; Jonathan Mitchell for maintenance and animal care at the UCR vivarium.

This research was supported by initial complement funding to PHD from UIUC and UCR and DC was supported by the Department of Microbiology from UIUC. LL was supported by the National Science Foundation Graduate Research Fellowship. Gnotobiotic mouse work and AH were supported by National Institute of General Medical Sciences grant R35GM124724. RJW is supported by the Allen Foundation with an Allen Distinguished Investigator Award.

## SUPPLEMENTARY FIGURE LEGENDS

**Figure S1. Transcriptional activity of the BV01 prophage and surrounding chromosome.** RNAseq reads from wild-type lysogen (WT) and cured lysogen (ΔBV01) strains were mapped to the region, and coverage was normalized to the total number of reads mapping to the genome. One representative of three replicates shown for each. The average normalized read coverage for each genome is displayed as the y-axis maximum (grey line). Maximum read coverage for the region is indicated on the y-axis. Forward reads (red) and reverse reads (blue) were plotted separately. Locations of two putative BV01-encoded transcriptional regulators are indicated (BVU_RS14235, BVU_RS14475).

**Figure S2. Prevalence of prophage insertion adjacent to *tspO* in human gut samples.** Reads from 256 healthy human gut metagenomes were obtained from the Human Microbiome Project Healthy Human Subjects Study. (A) Reads were first mapped to representative sequences of *tspO* from *B. vulgatus* and *B. dorei*. Percent abundance *tspO* reads was calculated on a per sample basis as the number of reads mapping to *tspO* divided by the total number of reads. Histogram shows counts of samples. (B) Reads mapping to *tspO* were filtered to only include reads antisense to *tspO*, predicted to point toward the *attB* based on the known genomic architecture. Mates to those reads were subsequently mapped to BV01 and its *Salyersvirus* relatives (Fig. 6). Only samples with read pairs bridging *tspO* and a phage sequence are shown (*n*=34). Percent *tspO* reads mated with *Salyersvirus* reads was calculated as the read pairs bridging *tspO* and a phage sequence divided by the total number of reads mapping antisense to *tspO*. Histogram shows counts of samples.

**Figure S3. Paired whole genome tree and nucleotide alignment of *Salyersviridae* phages.** Phylogenomic Genome-BLAST Distance Phylogeny implemented with the VCTOR online tool (50) using amino acid data from all phage ORFs. For consistency, all phage genomes were annotated with MetaGeneAnnotator (94) implemented via VirSorter (95). Support values above branches are GBDP pseudo-bootstrap values from 100 replications. Genome alignment of all phages made with MAUVE. One locally collinear block (LCB) connects phages B124-14 and B40-8 to the *Salyersviridae* at the nucleotide level (green). Other LCB connecting lines removed for clarity.

**Figure S4. Confirmation of activity of three additional Salyersviruses.** DNase-treated culture supernatants for predicted Salyersviruses BV02 (A), BV04 (B), and BV05 (C) were sequenced. Assembly resulted in contigs corresponding to the free form of phages BV02 and BV05; BV04 did not yield any contigs corresponding to the putative prophage region, suggesting it is inactivated. Assembled free phage contigs were aligned to their integrated prophage region with Mauve (A-C). Sequence reads were mapped back to their free and integrated forms and represented as coverage curves (A-C). Vertical orange lines indicate the location of *att* sequences; vertical dashed blue lines indicate the location of contig breaks. (D) PCR amplification with phage-specific primers tests for phage presence in pellet and supernatant fractions for 7 predicted Salyersviruses. Supernatant fractions were treated with DNase, eliminating all contaminating host genomic DNA, as demonstrated by the amplification of a host marker gene (16S rRNA). BV04 is not detectable in supernatant, supporting the conclusion that it is an inactivated prophage. PCR amplicons were visualized by agarose gel electrophoresis alongside GeneRuler Express DNA ladder (16S rRNA); ladder band sizes shown in Kb.

**Figure S5. *Salyersviridae* sequence is detectable in human-associated samples.** (A) Wastewater viromes were collected, and processed in three ways prior to sequencing (see Methods). Resulting reads were trimmed, pooled, and mapped to all *Salyersviridae* genomes and crAssphage. Only *Salyersvirinae* genomes shown in alignment to better demonstrate conservation, constructed with Mauve. Read coverages normalized to total number of reads in the metavirome. The maximum normalized read coverage for BC01 is 4.1, B40-8 and B124-14 is 1.13, and crAssphage is 7.33. (B) Reads from 256 healthy human gut metagenomes were obtained from the Human Microbiome Project Healthy Human Subjects Study. Reads were mapped to all 20 temperate *Salyersviridae* phages and crAssphage. Percent reads mapping was calculated on a per sample basis as the number of reads mapping to any virus divided by the total number of reads. Histogram shows counts of samples.

## SUPPLEMENTARY TABLE TITLES

Table S1. Rockhopper results for RNAseq from B. vulgatus WT (602), ΔBV01 (853) and ΔBV01ΔtspO (1662).

Table S2. Detection of *Salyersviridae* in Human Microbiome Project Healthy Human Subjects Study samples.

Table S3. Bacterial strains, plasmids, and primers used in this study.

Table S4. Breseq read mapping results of *B. vulgatus* WT and mutant strains used in this study.

Table S5. DNA sequencing data generated for B. vulgatus isolates, cell-free Salyersviridae phages, and wastewater viromes used in this study.

