## Supplemental Figures S1-S3 for "Infection with novel *Bacteroides phage BV01* alters host transcriptome and bile acid metabolism in a common human gut microbe"

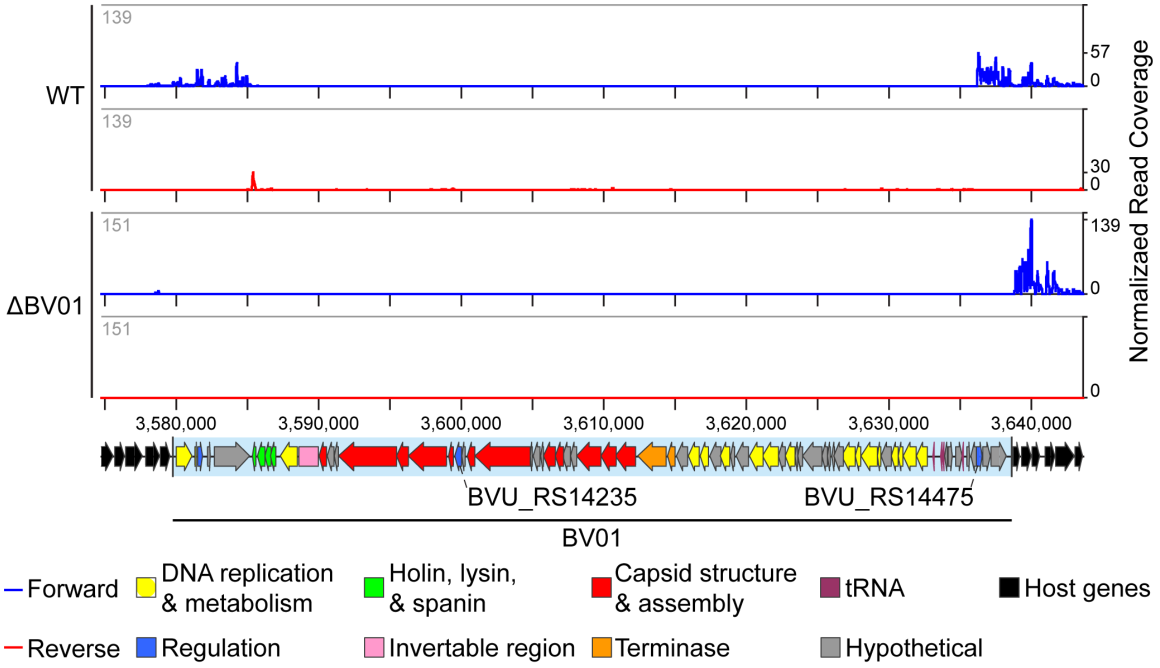


**Figure S1. Transcriptional activity of the BV01 prophage and surrounding chromosome.** RNAseq reads from wild-type lysogen (WT) and cured lysogen (ΔBV01) strains were mapped to the region, and coverage was normalized to the total number of reads mapping to the genome. One representative of three replicates shown for each. The average normalized read coverage for each genome is displayed as the y-axis maximum (grey line). Maximum read coverage for the region is indicated on the y-axis. Forward reads (red) and reverse reads (blue) were plotted separately. Locations of two putative BV01-encoded transcriptional regulators are indicated (BVU_RS14235, BVU_RS14475).

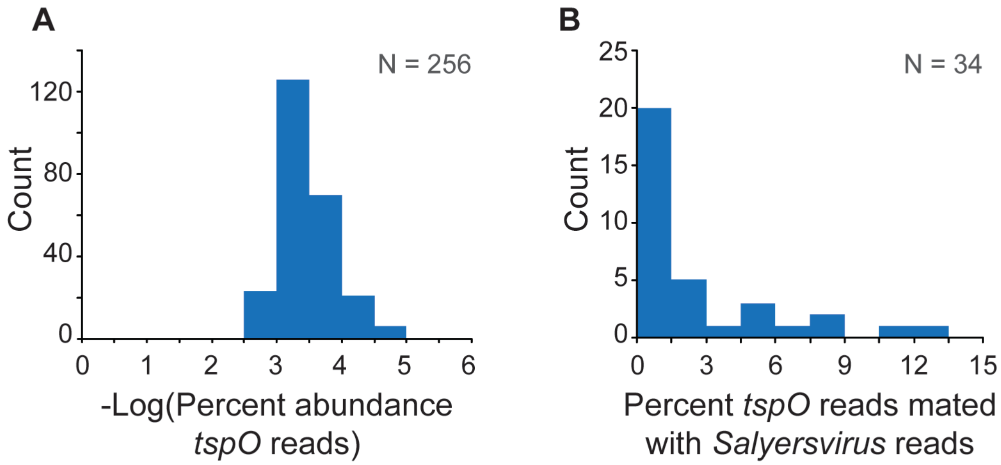


**Figure S2. Prevalence of prophage insertion adjacent to *tspO* in human gut samples.** Reads from 256 healthy human gut metagenomes were obtained from the Human Microbiome Project Healthy Human Subjects Study. (A) Reads were first mapped to representative sequences of *tspO* from *B. vulgatus* and *B. dorei*. Percent abundance *tspO* reads was calculated on a per sample basis as the number of reads mapping to *tspO* divided by the total number of reads. Histogram shows counts of samples. (B) Reads mapping to *tspO* were filtered to only include reads antisense to *tspO*, predicted to point toward the *attB* based on the known genomic architecture. Mates to those reads were subsequently mapped to BV01 and its *Salyersvirus* relatives (Fig. 6). Only samples with read pairs bridging *tspO* and a phage sequence are shown (*n*=34). Percent *tspO* reads mated with *Salyersvirus* reads was calculated as the read pairs bridging *tspO* and a phage sequence divided by the total number of reads mapping antisense to *tspO*. Histogram shows counts of samples.

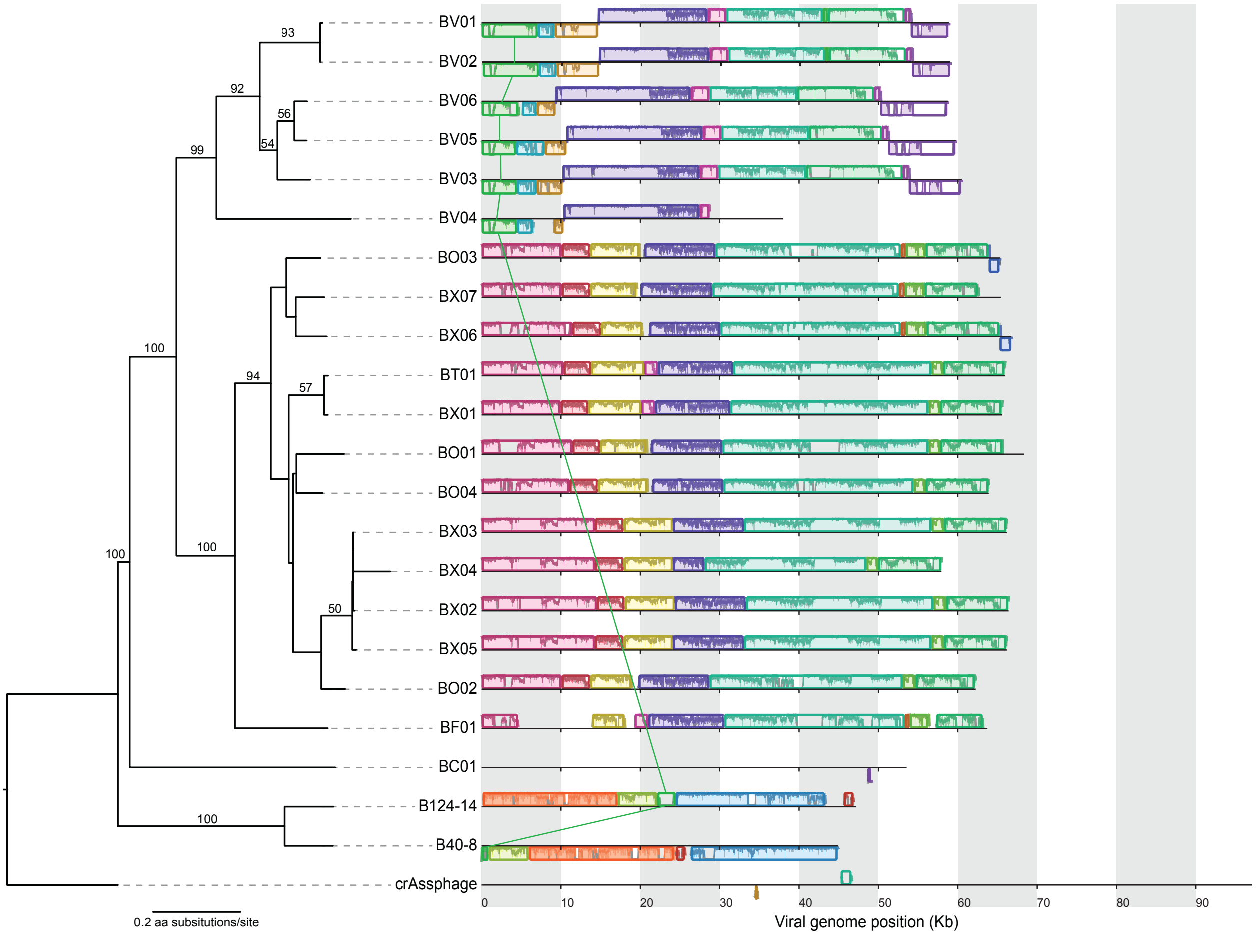


**Figure S3. Paired whole genome tree and nucleotide alignment of *Salyersviridae* phages.** Phylogenomic Genome-BLAST Distance Phylogeny implemented with the VCTOR online tool (53) using amino acid data from all phage ORFs. For consistency, all phage genomes were annotated with MetaGeneAnnotator (54) implemented via VirSorter (55). Support values above branches are GBDP pseudo-bootstrap values from 100 replications. Genome alignment of all phages made with MAUVE. One locally collinear block (LCB) connects phages B124-14 and B40-8 to the *Salyersviridae* at the nucleotide level (green). Other LCB connecting lines removed for clarity.
