## Supplemental Figures S4, S5 for "Infection with novel *Bacteroides phage BV01* alters host transcriptome and bile acid metabolism in a common human gut microbe"

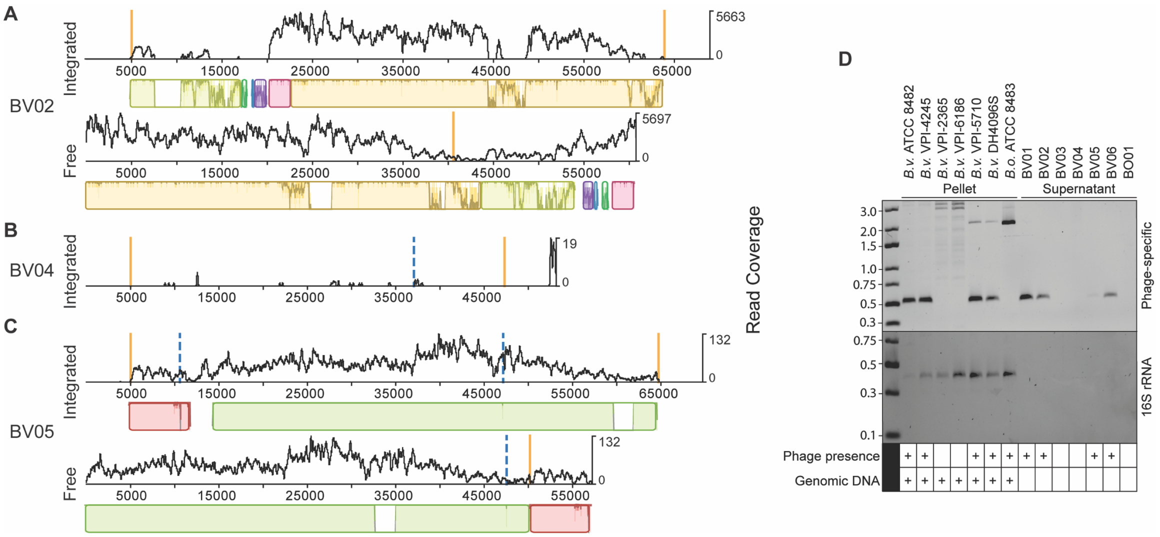


**Figure S4. Confirmation of activity of three additional Salyersviruses.** DNase-treated culture supernatants for predicted Salyersviruses BV02 (A), BV04 (B), and BV05 (C) were sequenced. Assembly resulted in contigs corresponding to the free form of phages BV02 and BV05; BV04 did not yield any contigs corresponding to the putative prophage region, suggesting it is inactivated. Assembled free phage contigs were aligned to their integrated prophage region with Mauve (A-C). Sequence reads were mapped back to their free and integrated forms and represented as coverage curves (A-C). Vertical orange lines indicate the location of *att* sequences; vertical dashed blue lines indicate the location of contig breaks. (D) PCR amplification with phage-specific primers tests for phage presence in pellet and supernatant fractions for 7 predicted Salyersviruses. Supernatant fractions were treated with DNase, eliminating all contaminating host genomic DNA, as demonstrated by the amplification of a host marker gene (16S rRNA). BV04 is not detectable in supernatant, supporting the conclusion that it is an inactivated prophage. PCR amplicons were visualized by agarose gel electrophoresis alongside GeneRuler Express DNA ladder (16S rRNA); ladder band sizes shown in Kb.

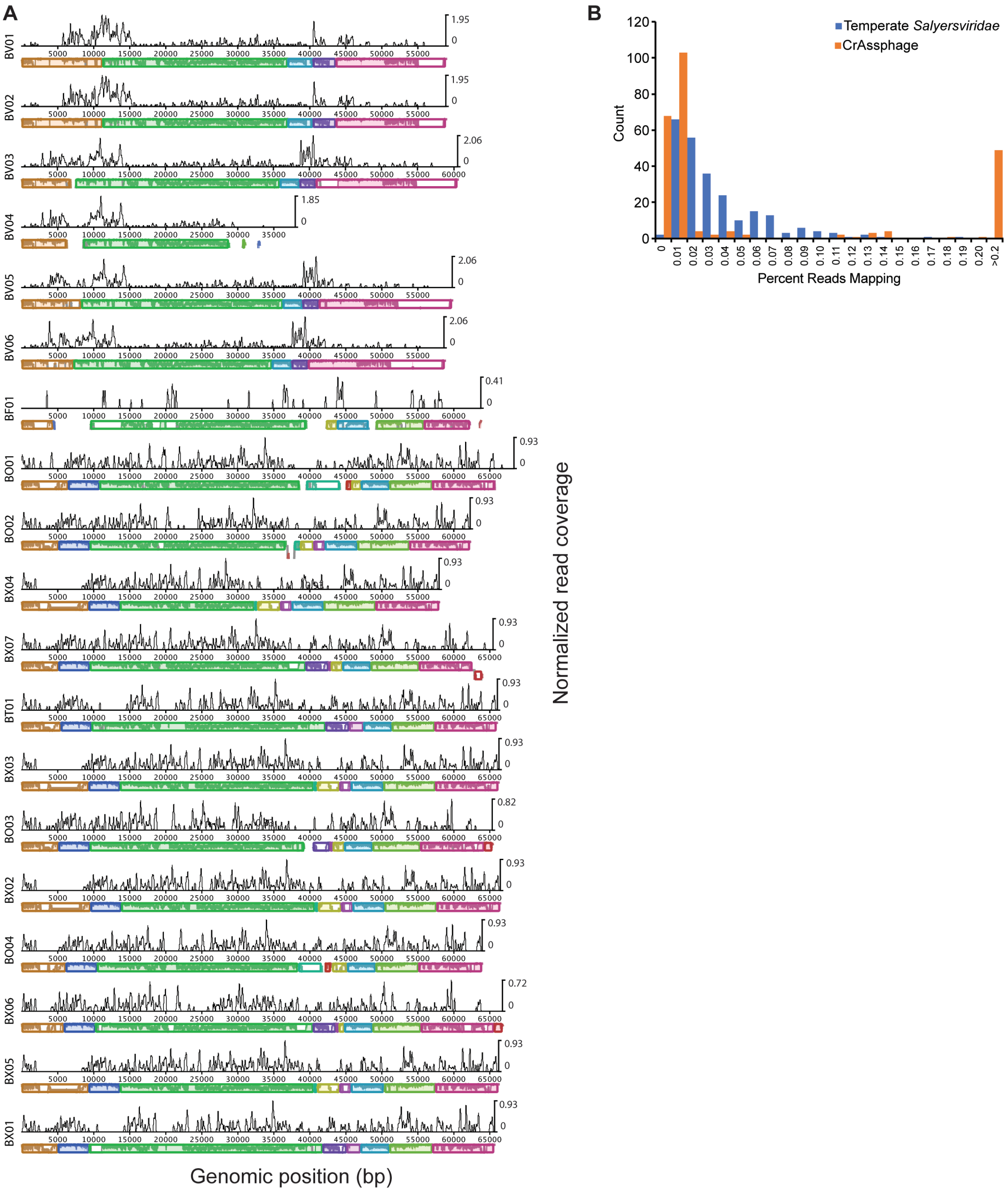


**Figure S5. *Salyersviridae* sequence is detectable in human-associated samples.** (A) Wastewater viromes were collected, and processed in three ways prior to sequencing (see Methods). Resulting reads were trimmed, pooled, and mapped to all *Salyersviridae* genomes and crAssphage. Only

*Salyersvirinae* genomes shown in alignment to better demonstrate conservation, constructed with Mauve. Read coverages normalized to total number of reads in the metavirome. The maximum normalized read coverage for BC01 is 4.1, B40-8 and B124-14 is 1.13, and crAssphage is 7.33. (B) Reads from 256 healthy human gut metagenomes were obtained from the Human Microbiome Project Healthy Human Subjects Study. Reads were mapped to all 20 temperate *Salyersviridae* phages and crAssphage. Percent reads mapping was calculated on a per sample basis as the number of reads mapping to any virus divided by the total number of reads. Histogram shows counts of samples.
